# A Pan-Pneumovirus vaccine based on immunodominant epitopes of the fusion protein

**DOI:** 10.1101/2022.02.24.481832

**Authors:** Jiachen Huang, Rose Miller, Jarrod Mousa

**Affiliations:** Department of Infectious Diseases, College of Veterinary Medicine, University of Georgia, Athens, GA 30602; Center for Vaccines and Immunology, College of Veterinary Medicine, University of Georgia, Athens, GA 30602; Department of Biochemistry and Molecular Biology, Franklin College of Arts and Sciences, University of Georgia, Athens, GA

## Abstract

Respiratory syncytial virus (RSV) and human metapneumovirus (hMPV) are two leading causes of severe respiratory infections in children, the elderly, and immunocompromised patients. The fusion (F) protein is the major target of neutralizing antibodies. Recent developments in stabilizing the pre-fusion conformation of the F proteins, and identifying immunodominant epitopes that elicit potent neutralizing antibodies have led to testing of numerous pre-fusion RSV F-based vaccines in clinical trials. We designed and tested the immunogenicity and protective efficacy of a chimeric fusion protein that contains immunodominant epitopes of RSV F and hMPV F (RHMS-1). RHMS-1 has several advantages over vaccination with pre-fusion RSV F or hMPV F, including a focus on recalling B cells to the most important protective epitopes and the ability to induce protection against two viruses with a single antigen. RHMS-1 was generated as a trimeric recombinant protein, and negative-stain EM analysis demonstrated the protein resembles the pre-fusion conformation. Probing of RHMS-1 antigenicity using a panel of RSV and hMPV F-specific monoclonal antibodies (mAbs) revealed the protein retains features of both viruses, including the pre-fusion site Ø epitope of RSV F. BALB/c mice immunized with RHMS-1 had serum binding and neutralizing antibodies to both viruses. RHMS-1 vaccinated mice challenged with RSV or hMPV had undetectable virus in lung homogenates for both viruses, in contrast to RSV F or hMPV F vaccinated mice, which had detectable virus for hMPV and RSV, respectively. Overall, this study demonstrates protection against two viruses with a single antigen and supports further testing of RHMS-1 in additional pre-clinical animal models.

**Significance:** Respiratory syncytial virus (RSV) and human metapneumovirus (hMPV) are two Pneumoviruses that cause substantial respiratory tract infections in young children, the elderly, and immunocompromised adults. Yet vaccines against either of them are still unavailable. Multiple promising vaccine designs based on the fusion proteins of RSV and hMPV have been tested, but none of them can induce cross-protective antibody responses despite their similar structures and closely related amino acid sequence identity. Here, we report a vaccine candidate, RHMS-1, which combines the immunodominant epitopes of the RSV and hMPV fusion proteins into a single antigen. RHMS-1 maintains the immunological features of both RSV F and hMPV F, which can be recognized by epitope-specific mAbs and human B cells pre-exposed to RSV or hMPV. Furthermore, this is the first immunogen that induced potent cross-neutralizing antibodies and protected mice from RSV and hMPV challenge. Our results suggest RHMS-1 is a promising Pan-Pneumovirus vaccine.

## Introduction

Respiratory syncytial virus (RSV) and human metapneumovirus (hMPV) are significant causes of acute lower respiratory tract infections (ALRI) in infants and young children (1–4). RSV was first identified in 1956, and was subsequently recognized as a common cause of respiratory illness in early life (5). The majority of children experience at least one RSV infection before 2 years of age, and infants under 6 months old have a higher risk of severe disease requiring hospitalization (6). hMPV was identified in 2001, and it has been found to be the second most common cause of viral lower respiratory infection in children (7). In contrast to RSV, the peak age for infant hospitalizations caused by hMPV infections is 6-12 months old, with a nearly 100% exposure rate by the age of 5 (3). Reinfections of both RSV and hMPV are common throughout life, which usually cause mild symptoms in healthy adults. However, for certain populations including immunocompromised patients, individuals over 65 years of age, and people with underlying conditions such asthma or chronic obstructive pulmonary disease (COPD), infection with RSV or hMPV may lead to severe bronchiolitis and pneumonia (8–11).

The fusion (F) glycoproteins of RSV and hMPV are highly similar in structure and share ∼30% amino acid sequence identity. Both F proteins belong to the class I viral fusion protein family and play indispensable roles in viral attachment as well as membrane fusion. To become fusion competent, the F0 precursor must to be cleaved into F1 and F2 subunits that are linked by two disulfide bonds to generate a mature meta-stable homotrimer (12). RSV F is cleaved at two furin cleavage sites with the p27 fragment in between F1 and F2 removed, whereas hMPV F has only one cleavage site that can be cleaved by the host membrane protease TMPRSS2 (13). To initiate the fusion process, the hydrophobic fusion peptide on the N terminus of the F2 subunit is exposed and inserted into host cell membrane, which triggers the conformational rearrangements that turn the F protein into the stable post-fusion state, and brings the viral and host cell membranes together for lipid mixing.

Multiple antigenic sites have been identified on both RSV F and hMPV F proteins. Among the six known antigenic sites of RSV F, pre-fusion-specific sites Ø and V are targeted by over 60% of neutralizing antibodies in humans (14, 15), indicating these sites are vital for immune recognition and antibody neutralization (16). hMPV F shares three antigenic sites (III, IV, V) with RSV F, as several antibodies have been found to cross react with RSV and hMPV F at these epitopes (17–20). In addition, the area in between site III and IV was found to be a distinct hMPV site that is recognized by a mAb called DS7 (21, 22). Studies have shown that, unlike RSV F-specific antibodies, the majority of hMPV F-specific antibodies target epitopes present in both pre-fusion and post-fusion conformations, likely due to glycosylation present near pre-fusion-specific sites on the head of hMPV F (23, 24).

Currently, there are no vaccines available for either RSV or hMPV. Previous attempts with formalin-inactivated RSV and hMPV vaccines revealed that low affinity, non-neutralizing F-specific antibodies induced by denatured fusion proteins cannot provide protection, and lead to vaccine enhanced disease (25–29). By stabilizing the F proteins in the pre-fusion conformation, several studies have demonstrated improvement in neutralizing antibody titers (30, 31), while for hMPV, pre-fusion and post-fusion F proteins induced comparable neutralizing antibodies in mice (24) and immunization with post-fusion F completely protected mice from hMPV challenge (32). Several epitope-focused vaccine designs have been tested for RSV and hMPV F. A head-only RSV F protein boosted titers of neutralizing Abs targeting antigenic sites Ø and II (33). In a different study, RSV F was modified by glycan-masking that blocked poorly neutralizing epitopes on a nanoparticle, which induced a more potent neutralizing Ab response than a prefusion F trimer (34). Based on computational protein design strategies, RSV F site II was presented on a scaffold fused with RSV N-based nanoparticles, which boosted subdominant neutralizing antibody responses targeting antigenic site II in mice (35, 36). In addition, RSV F neutralizing antigenic sites (Ø, II, IV) were tested on de novo protein scaffolds respectively, and a mixture of these epitope-based immunogens induced focused immune responses toward the target antigenic sites (37). All of the studies above demonstrate the concept that engineered RSV F epitope-based immunogens can induce and boost neutralizing RSV F antibodies.

The idea of universal vaccine development provides the possibility of preventing multiple variants (like flu and HIV) by a single immunogen. Due to the similarities between RSV and hMPV F, researchers have tried to generate universal RSV/MPV vaccines by grafting the helix-turn-helix motif (site II) from RSV F onto hMPV F, however, this chimera induced neutralizing antibody responses only to hMPV, but not RSV (38). A similar study that grafted RSV F and hMPV F epitopes on pre-fusion and pos-fusion F proteins showed that chimeric proteins swapping either site II or site IV can induce cross-neutralizing antibodies in mice, but a challenge with pre-fusion candidates was lacking (39). For the influenza hemagglutinin (HA) protein, chimeric immunogens generated by swapping the HA head with zoonotic subtypes while retaining the conserved HA stem successfully induced antibodies targeting the subdominant HA stem (40, 41).

Based on these findings and the knowledge about structures and the immunodominant epitopes of RSV and hMPV F proteins, we designed a novel chimeric immunogen that contains the head of RSV F and the stem of hMPV F. The **R**SV **h**ead h**M**PV **s**tem construct **1** (RHMS-1) protein was stably expressed as a pre-fusion trimer that preserved the structural features on key antigenic sites for both RSV and hMPV F proteins. RHMS-1 retains immunodominant epitopes of both F proteins, including antigenic sites Ø and V of RSV F, and sites IV, DS7, and III of hMPV F. Immunization of mice with RHMS-1 induced potent neutralizing antibodies that protected mice from both RSV and hMPV challenge. Overall, our data demonstrates that RHMS-1 can be a promising universal vaccine against both Pneumoviruses.

## Material and methods

### Ethics statement

All animal studies were approved by the University of Georgia Institutional Animal Care and Use Committee. Human studies were approved by the University of Georgia Institutional Review Board, and subjects were recruited to the University of Georgia Clinical and Translational Research Unit and written informed consent was obtained.

### Expression and purification of proteins

Plasmids encoding cDNAs of Pneumovirus proteins were synthesized (GenScript) and cloned into the pcDNA3.1^+^ vector (30, 32, 42). The stable cell line that expresses the hMPV B2 F protein was utilized as previously described (32). The rest of the F proteins and monoclonal antibodies were transiently expressed in Expi293F cells. The proteins were harvested from the supernatant of cell cultures and purified by HisTrap Excel (for his-tagged proteins) or Protein G (for antibodies) columns (GE Healthcare Life Sciences). RHMS-1 (trimer), RSV A2 F (trimer), and trypsin-treated hMPV B2 F (monomer) were further purified by size exclusion chromatography on a Superdex S200, 16/600 column (GE Healthcare Life Sciences).

### Negative-stain electron microscopy analysis

Purified RHMS-1 (trimer) was applied on carbon-coated copper grids (5 μl of 10 μg/mL protein solution) for 3 min. The grid was washed in water twice then stained with Nano-W (Nanoprobes) for 1 min. Negative-stain electron micrographs were acquired using a JEOL JEM1011 transmission electron microscope equipped with a high-contrast 2K-by-2K AMT midmount digital camera.

### ELISA of RHMS-1 with mAbs or human/mouse serum

384-well plates (Greiner Bio-One) were coated with 2 μg/ml of antigen in PBS overnight at 4°C. The plates were then washed once with water before blocking for 1 hour with blocking buffer. Primary mAbs (starting at 20 μg/ml and followed by 3-fold dilutions) or serial dilutions of human/mouse serum (starting with 1:50 and followed by 3-fold dilutions) were applied to wells for 1 hour after three washes with water. Plates were washed with water three times before applying 25 μl of secondary antibody (goat anti-human IgG Fc-Southern Biotech, 2048-04; goat anti-mouse IgG Fc-Southern Biotech, 1033-04) at a dilution of 1:4,000 in blocking buffer. After incubation for 1 hour, the plates were washed five times with 0.05% PBS-Tween-20, and 25 μl of a PNPP (p-nitrophenyl phosphate) substrate solution (1 mg/ml PNPP in 1 M Tris base) was added to each well. The plates were incubated at room temperature for 1 hour before reading the optical density at 405 nm (OD_405_) on a BioTek plate reader. Data were analyzed in GraphPad Prism using a nonlinear regression curve fit and the log(agonist)-versus-response function to calculate the binding EC_50_ values. Mouse serum IgG titers were calculated from the highest dilution of a serum sample that produced OD_405_ readings of >0.3 above the background readings and were shown in a log_10_ scale as previously described (32).

### ELISA screening human PBMCs

As previously described (43), peripheral blood mononuclear cells (PBMCs) and plasma were isolated from human subject blood samples using CPT tubes (BD, 362753), and PBMCs were frozen in the liquid nitrogen vapor phase until further use. For serology screening, the plasma of 41 subjects were used for ELISA as described above. The IgG binding was quantified by area under the curve (AUC) values using GraphPad Prism. For PBMC screening, ten million PBMCs were mixed with 8 million previously frozen and gamma irradiated NIH 3T3 cells modified to express human CD40L, human interleukin-21 (IL-21), and human B-cell activating factor (BAFF) in 80 mL StemCell medium A (StemCell Technologies) containing 1 μg/mL of cyclosporine A (Millipore-Sigma). The mixture of cells was plated in four 96-well plates at 200 μl per well in StemCell medium A. After 6 days, culture supernatants were screened by ELISA for IgG binding to the RHMS-1 (trimer), RSV A2 DsCav1 F (trimer), and trypsin-treated hMPV B2 F (monomer). Each well is represented by a dot with the OD_405_ against RSV/hMPV F as the x coordinate, and the OD_405_ against RHMS-1 as the y coordinate.

### Animal immunization and hMPV/RSV challenge

BALB/c mice (6 to 8 weeks old; The Jackson Laboratory) were immunized in a prime-boost regimen with purified RHMS-1 (trimer), RSV A2 DsCav1 F (trimer), or trypsin-treated hMPV B2 F (monomer) (20 μg protein/mouse) + an equal volume of AddaS03 adjuvant via the subcutaneous route into the loose skin over the neck, while mice in control groups were immunized with PBS + AddaS03 adjuvant. Three weeks after prime, the mice were boosted with the same amount of the antigens + adjuvant. Three weeks after the boost, mice were bled and then intranasally challenged with RSV A2 (2.8×10^6^ PFU per mouse) or hMPV TN/93-32 (3×10^5^ PFU per mouse). Mice were sacrificed 5 days post-challenge, and lungs were collected and homogenized for virus titration as previously described (32). Briefly, RSV challenged lung homogenates were plated on HEp-2 cells (EMEM+2% FBS) while hMPV challenged lung homogenates were plated on LLC-MK2 cells (EMEM + 5 μg/ml trypsin-EDTA and 100 μg/ml CaCl_2_) in 24 well plates. After 4-5 days, the cells were fixed with 10% neutral buffered formalin and the plaques of both viruses were immunostained with mAbs MPV364 (for hMPV) or 101F (for RSV). Plaques were counted by hand under a stereomicroscope.

### Virus neutralization assays with immunized mice serum

For serum neutralization assays, the serum of 4/8 mice were randomly picked from each group. Heat-inactivated mouse serum was serially diluted (starting at 1:25 and followed by 3-fold dilutions) and incubated 1:1 with a suspension of hMPV (CAN/97-83 and TN/93-32) or RSV (A2 and B) for 1 hour at room temperature. PBS was mixed with viruses as negative control. LLC-MK2 (for hMPV) or HEp-2 cells (for RSV) in 24-well plates were then inoculated with the serum-virus mixture (50 μL/well) for 1 hour and rocked at room temperature before adding the overlay. After 4-5 days, the plaques were stained described above. The percent neutralization was calculated by (PFU in control wells − PFU in serum wells)/PFU in control wells × 100%.

## Results

### RHMS-1 was expressed as pre-fusion trimer

We designed the RHMS-1 protein based on the pre-fusion structures of RSV F (5UDE) and hMPV F (5WB0) using ChimeraX (44), and a model of RHMS-1 based on these structures is shown in **Figure 1A**. The design of RHMS-1 maintains the signal peptide, two cleavage sites, DsCav1 mutations (S115C, S190F, V207L, S290C) (30), and the fusion peptide of RSV F. Part of the F2 N-terminus (residues 26-54) and the F1 C-terminus (residues 315-531) was replaced by the homologous hMPV F sequences, with two junctions located on β2 and β7 strands. Two glycosylation sites (RSV F-N70, hMPV F-N353) are retained in RHMS-1. A GCN4 trimerization domain and a hexa-histidine tag were appended to the F1 C-terminus **(Figure 1B)**. Intact RSV F sites II, V, and Ø and hMPV F sites IV, DS7 were adopted from the original sequence. For antigenic site III, part of the cross-protomer site III of RHMS-1 is on the RSV head, but most of it is originated from hMPV F **(Figure 1C)**.

**Figure 1.**
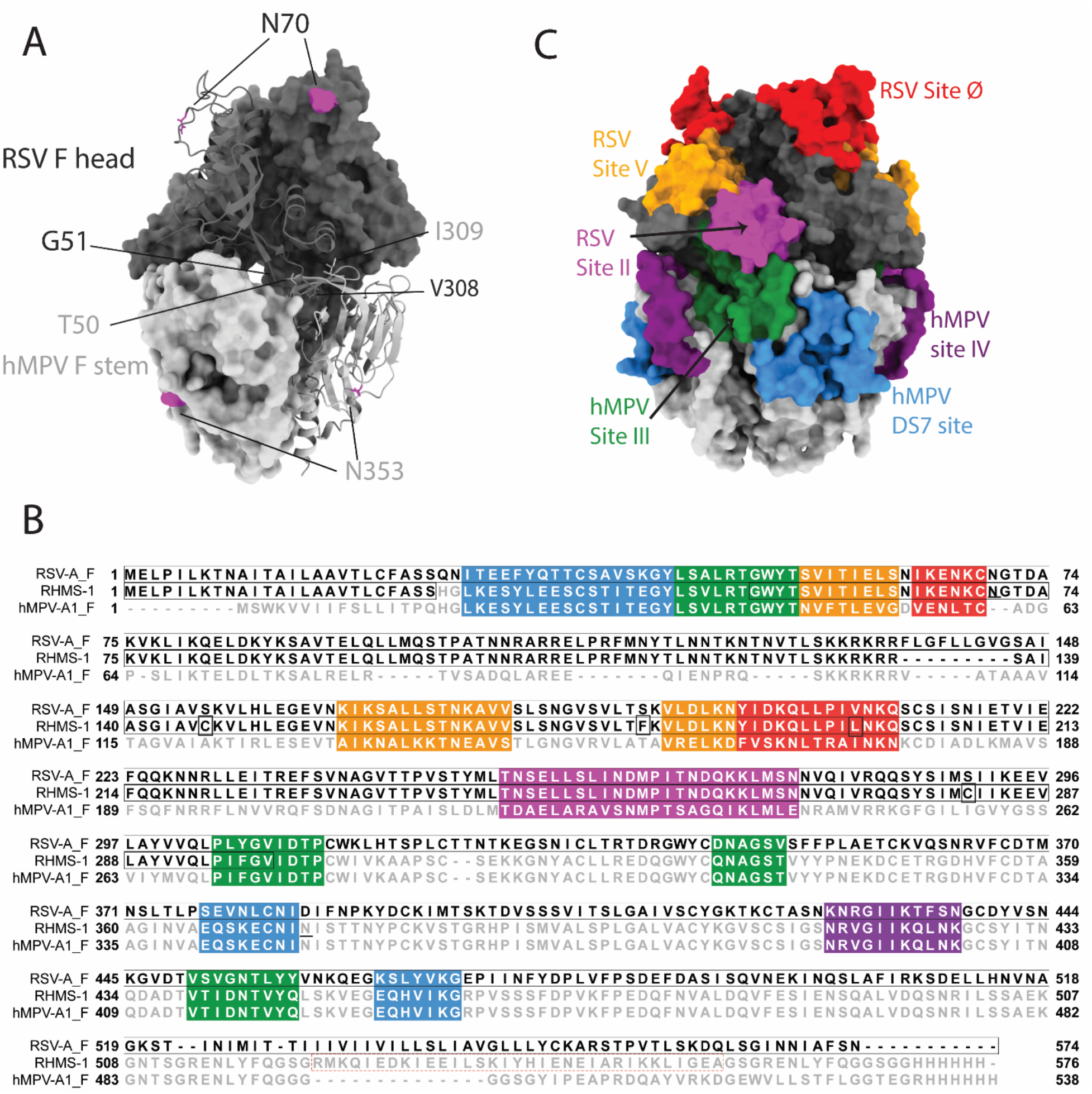
RHMS-1 protein design(A) The diagram generated with the head (gray) of pre-fusion RSV F (5UDE) and the stem (silver) of pre-fusion hMPV F (5WB0) shows one protomer in cartoon and the rest of two protomers in surface. (B) Sequence alignment of RSV-A F, RHMS-1, and hMPV-A1 F generated by Jalview. The sequences of known antigenic sites are highlighted: RSV site Ø – red, RSV site V – orange, RSV site II – magenta, hMPV site III – forest, hMPV site IV – purple, hMPV DS7 site – blue. In RHMS-1 sequence, four Ds-Cav1 mutations are circled in boxes, two N-linked glycosylation sites are underlined and the GCN4 trimerization domain is circled in red dashed box. (C) The antigenic sites colored in accordance with the sequences highlighted in Figure 1B are displayed on the diagram. Both figure 1A and C were made by ChimeraX.

RHMS-1 was expressed in HEK293F cells and size exclusion chromatography (SEC) showed RHMS-1 was mainly expressed as a trimeric protein, but the size is slightly bigger than RSV F trimers **(Figure 2A)**. For hMPV F, trimers and monomers were observed after trypsin treatment as previously described (32), but the size of trimeric hMPV F is smaller than RHMS-1 and RSV F, likely due to trypsinization **(Figure 2A)**. Like RSV F, RHMS-1 was deduced to be cleaved after expression, as the F2 domain was observed on the gel under reducing conditions **(Figure 2B)**, and the sizes of RHMS-1 and RSV F bands are consistent with the peaks shown in **Figure 2A**. Negative-stain EM analysis demonstrated that majority of the particles are in pre-fusion conformation based on “ball-like” structures resembling pre-fusion RSV and hMPV F (24, 30) **(Figure 2C)**, indicating the DsCav1 mutations work well in stabilizing the structure of RHMS-1.

**Figure 2.**
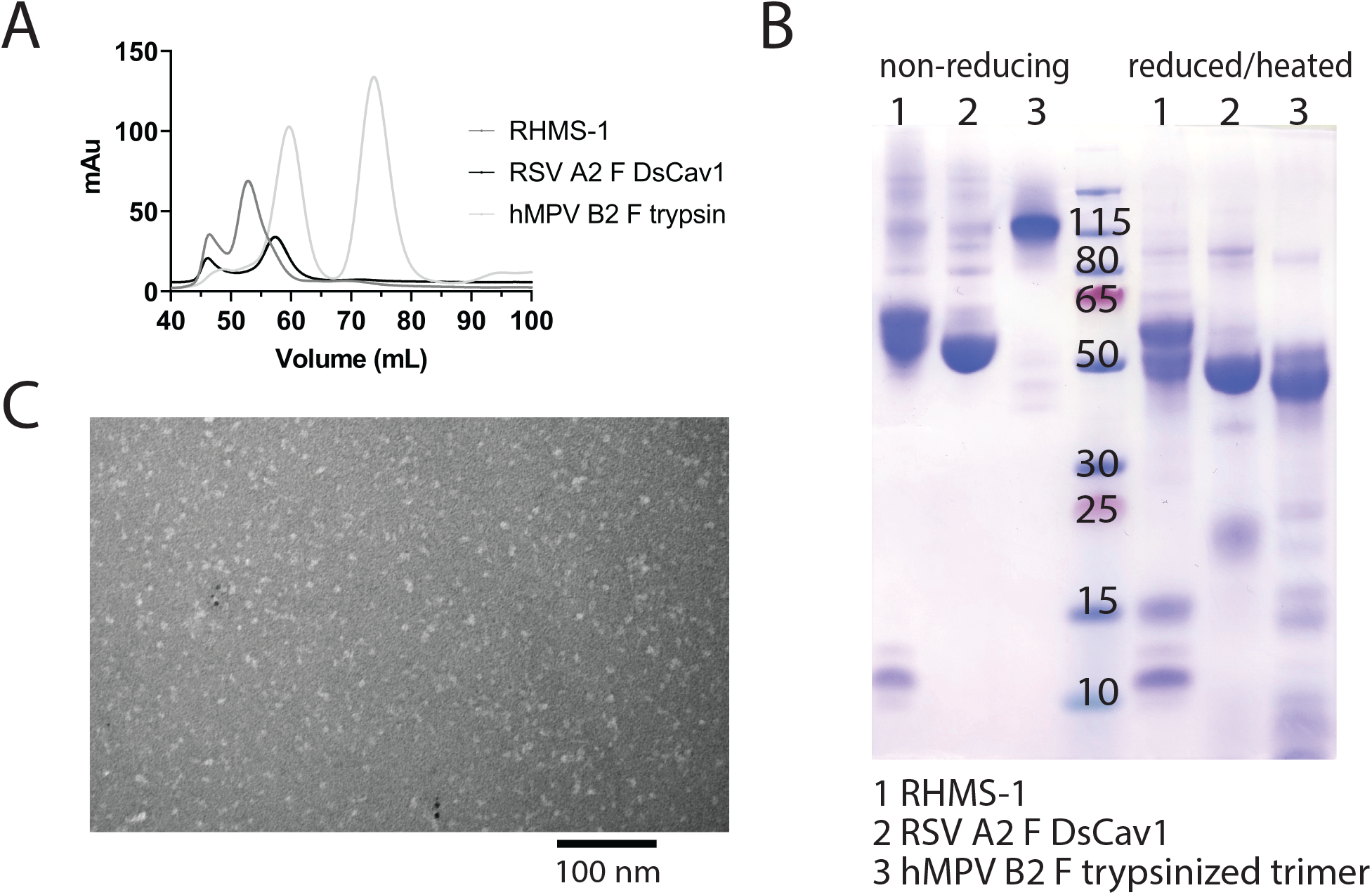
Purification and Negative-stain EM of RHMS-1. (A) Size exclusion chromatography curves of RHMS-1 (gray), RSV A2 F DsCav1 (black), and trypsinized hMPV B2 F (silver). (B) SDS-PAGE of F proteins in non-reducing and reduced/heated conditions. (C) Representative negative-stain electron micrograph of RHMS-1 obtained from fractions 50-60 mL from the size exclusion chromatogram shown in (A), scale bar: 100 nm.

### RHMS-1 shares immunological features of both RSV and hMPV F

To determine if RHMS-1 retains the correct conformation of each antigenic site, mAbs specifically targeting these sites were tested for binding by ELISA **(Figure 3A)**. For both RSV F and RHMS-1, mAb D25 binds to site Ø (45) and motavizumab binds to site II (46, 47) at similar EC_50_ values **(Figure 3B)**. For both hMPV F and RHMS-1, mAbs DS7 and MPV196 bind to the DS7 site (21, 43). mAb 101F binds to all three antigens on site IV (17) while mAb MPE8 binds to site III on RSV F and RHMS-1 (48), but not monomeric hMPV F, likely due to the cross-protomer epitope that is only partially displayed on a single monomer. The binding site of mAb MPV364 partially overlaps with hMPV site III, but it was also predicted to interact with the head of hMPV F (43), and mAb MPV458 binds to the 66-87 peptide on the head of hMPV F (42), therefore, both MPV364 and 458 do not bind to RHMS-1 as expected.

**Figure 3.**
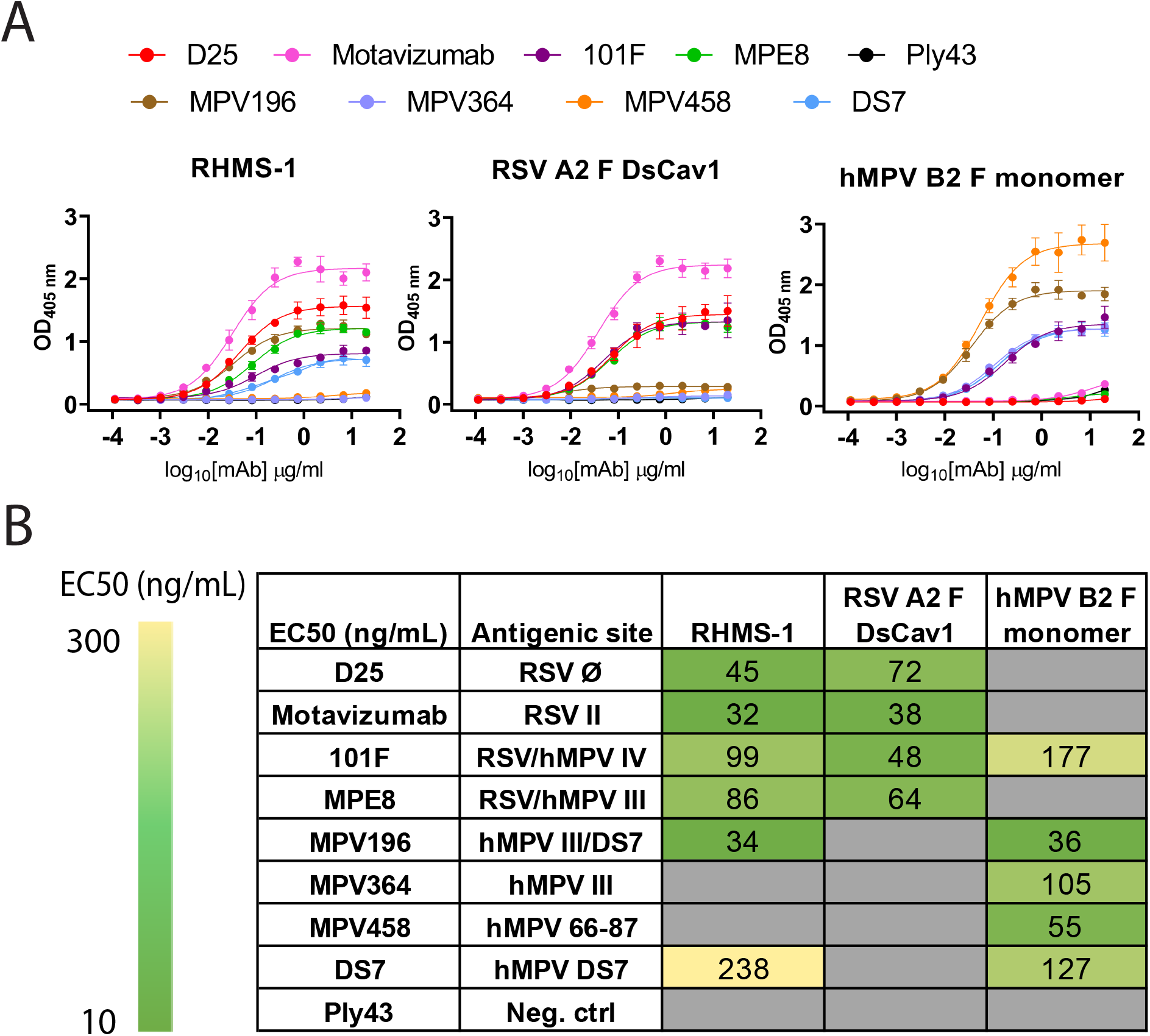
Antigenic site-specific mAbs binding to F proteins. (A) ELISA binding curves of mAbs targeting different RSV/hMPV F antigenic sites against RHMS-1, RSV A2 F DsCav1, and trypsinized hMPV B2 F monomer. (B) EC_50_ values of the binding curves in (A). The binding curves and the EC_50_ values were generated by GraphPad Prism.

### RHMS-1 can be recognized by B cells pre-exposed to RSV F or hMPV F

To verify if epitopes on RHMS-1 can be recognized by the human immune system in a similar manner comparing to RSV F or hMPV F, we screened plasma IgG responses from 41 human subjects against all 3 proteins by ELISA. Overall, positive correlations of serum IgG bindings were observed for both RSV F vs. RHMS-1 **(Figure 4A)** and hMPV F vs. RHMS-1 **(Figure 4B)**, suggesting epitopes are conserved between the native F proteins and the epitopes included on RHMS-1. The antibody responses at the cellular level were also tested by measuring the binding of supernatant from stimulated B cells in four subjects. The B cells in PBMCs were activated through coincubation with NIH 3T3 cells expressed human CD40L, human interleukin-21 (IL-21), and human B-cell activating factor (BAFF) to stimulate growth and IgG secretion to the culture supernatant as previously described (43). For all of the subjects we tested, the majority of RSV F-positive B cells are also positive for RHMS-1 **(Figure 5A)**. This finding is expected as the majority of human B cells target the head of the RSV F protein, which is retained in RHMS-1. Such correlations are still present for hMPV F **(Figure 5B)**, although the frequencies of hMPV F-positive B cells are generally lower than RSV F-positive B cells, therefore, we see populations of hMPV F negative, RHMS-1 positive B cells close to the Y axes.

**Figure 4.**
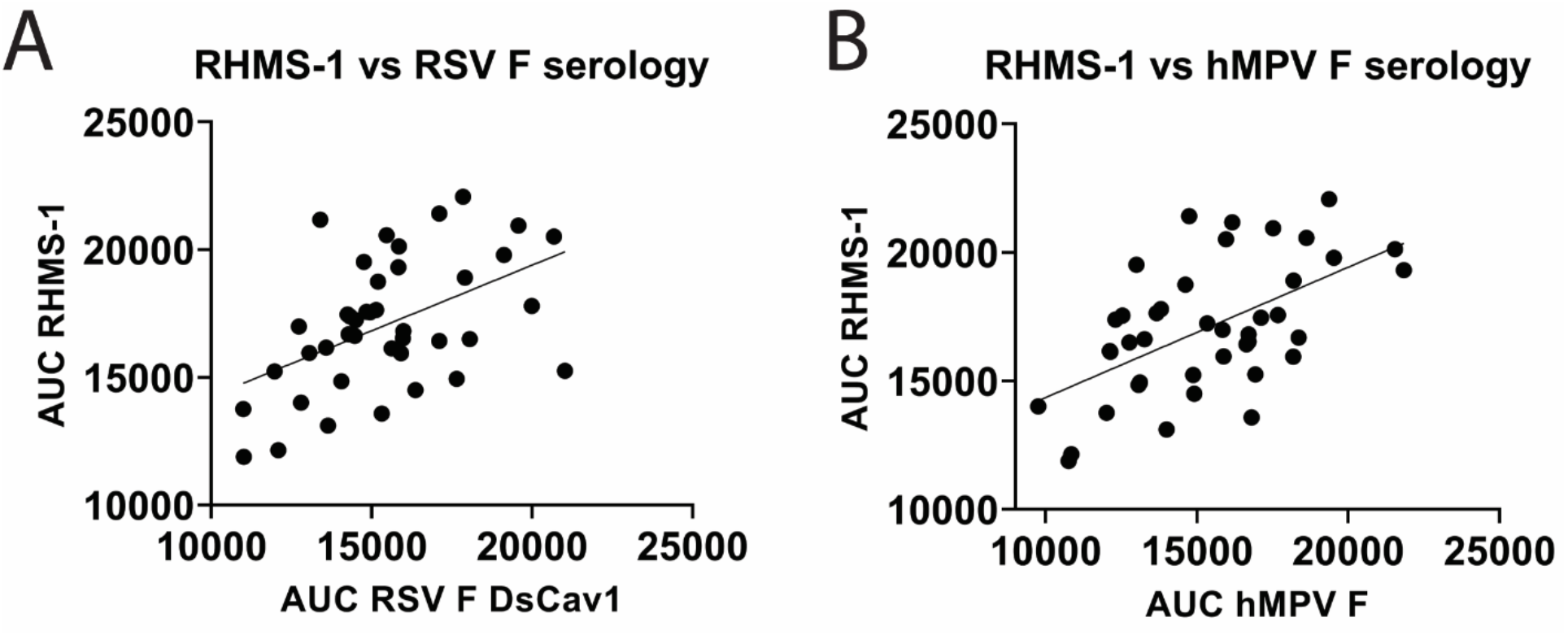
Serology of human plasma against F proteins. Area under the curve analysis of plasma IgG binding to RHMS-1 vs. RSV A2 F DsCav1 (A) and RHMS-1 vs. trypsinized hMPV B2 F monomer (B). Each dot represents one subject, and the lines indicate the linear regression fit of the data sets. Figures and data analysis was generated by GraphPad Prism.

**Figure 5.**
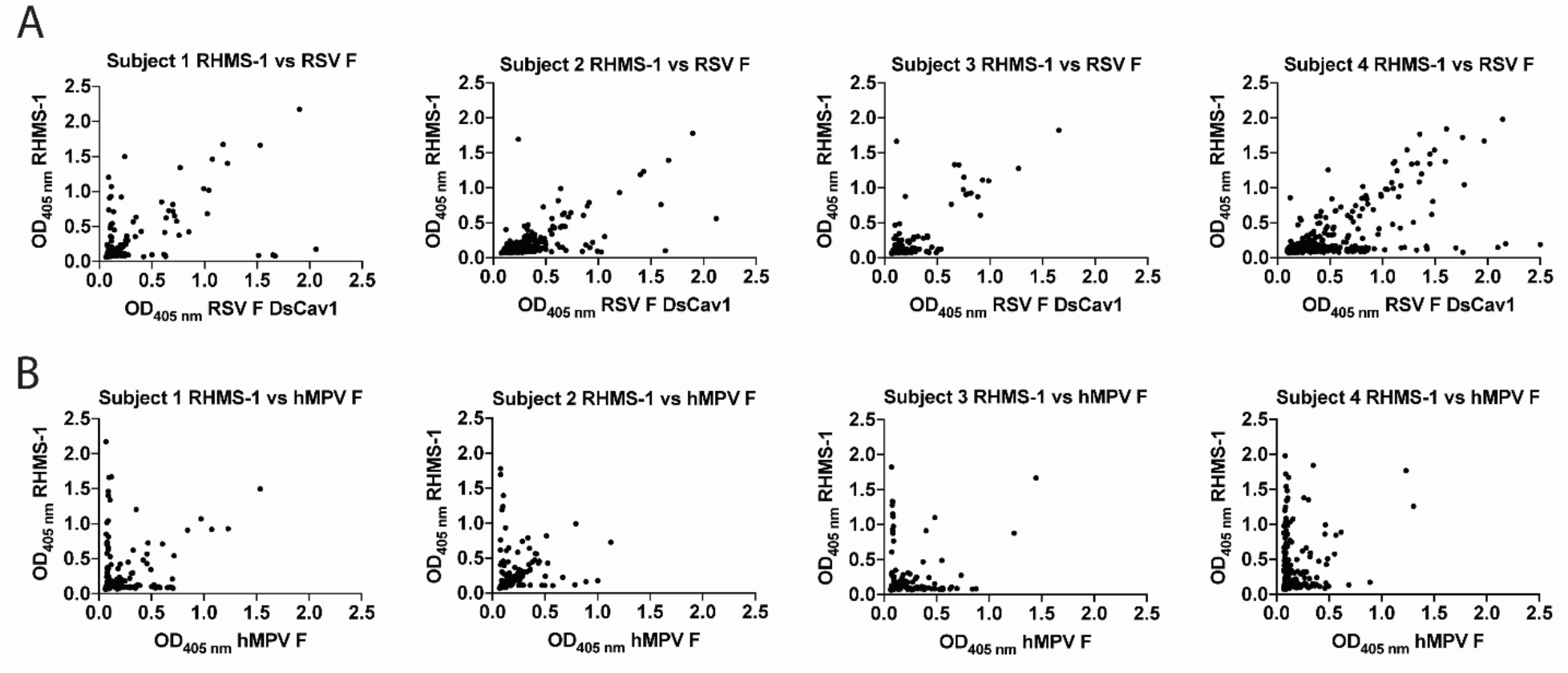
Human PBMCs binding to F proteins. ELISA OD_405 nm_ values of B cell culture supernatants binding to RHMS-1 vs. RSV A2 F DsCav1 (A) and RHMS-1 vs. trypsinized hMPV B2 F monomer (B). Each dot represents the B cell supernatant in a single well of a 384 well plate initially containing 20,000 PBMCs. Figures were generated by GraphPad Prism.

### Vaccination with RHMS-1 elicits potent neutralizing antibodies and cross-protection in mice

To evaluate the immunological properties of RHMS-1, it was tested as a vaccine in the mouse model. BALB/c mice were subcutaneously primed and boosted with 20 μg of RHMS-1, RSV F DsCav1, hMPV monomeric B2 F, or PBS in an emulsion formulated with AddaS03 adjuvant and then challenged with RSV or hMPV **(Figure 6A)**. All RHMS-1 vaccinated mice showed serum IgG binding titers against both RSV F DsCav1 and hMPV monomeric B2 F proteins **(Figure 6B, 6C)**. Representative viruses from each RSV subgroup and each hMPV genotype were neutralized by RHMS-1 immunized mouse serum 3 weeks after the boost **(Figure 6D & 6E)**. RSV F DsCav1 immunization failed to induce IgG that cross-recognize hMPV monomeric B2 F, while hMPV monomeric B2 F immunized mice showed moderate binding, but non-neutralizing IgG against RSV. Three weeks after the boost, mice were intranasally challenged RSV A2 or hMPV TN/93-32. The virus titers in the lung homogenate were determined 5 days post challenge. Vaccination with RHMS-1 completely protected the mice from challenges of both viruses, while RSV A2 F DsCav1 and hMPV B2 F monomer vaccinated groups protected mice only against the autologous virus **(Figure 6F)**.

**Figure 6.**
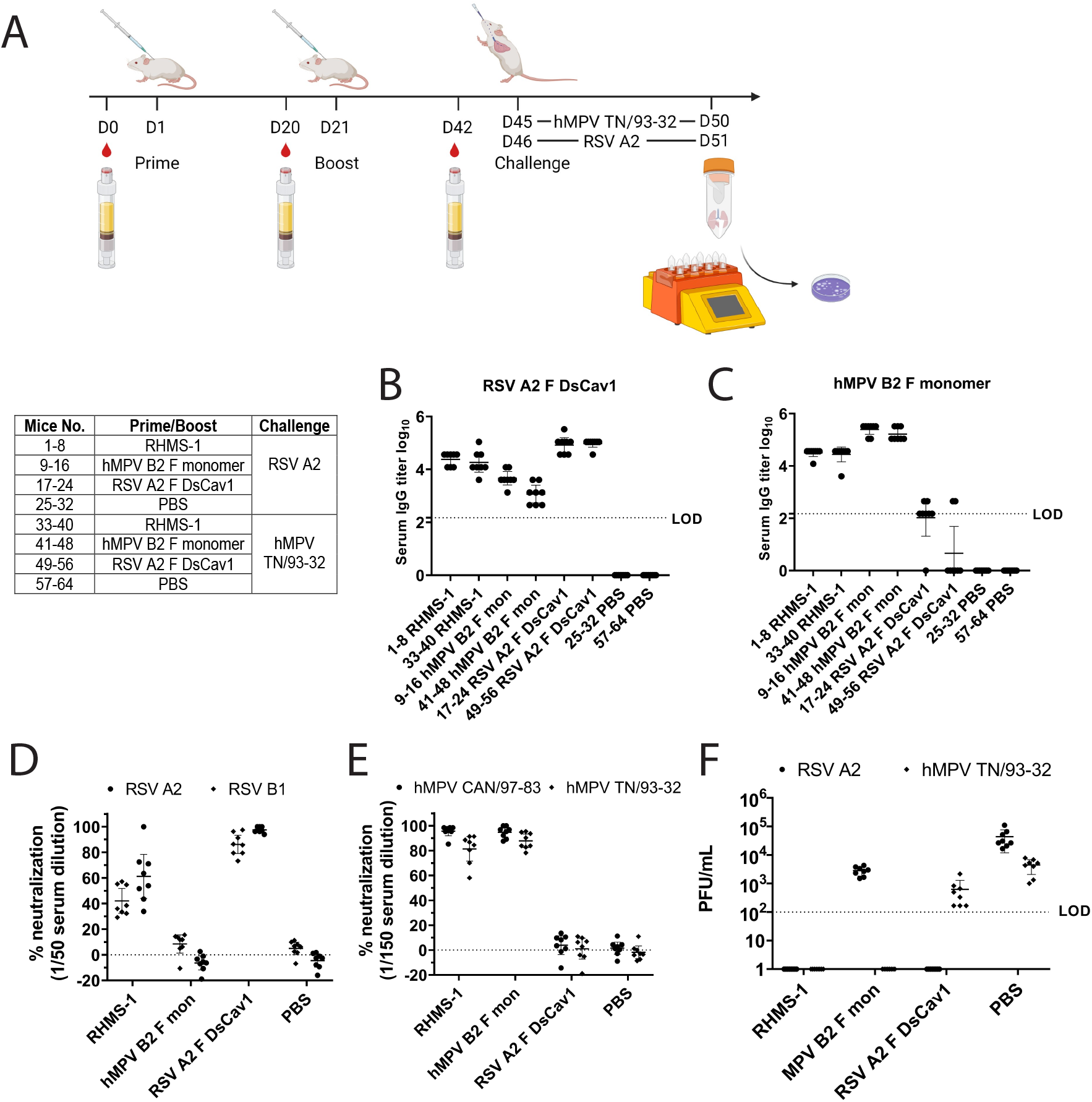
Mouse immunization and challenge studies. (A) Study regimen. (B) Day 42 serum IgG titers against RSV A2 F DsCav1, and (C) against trypsinized hMPV B2 F monomer before challenge. (D) Serum neutralization against RSV A2 and B1 strains at 1/50 dilution and against (E) hMPV CAN/97-83 and TN/93-32 at a 1/150 dilution. (F) Lung viral titers for RSV A2 and hMPV TN/93-32 in the lung homogenates of mice 5 days post-challenge. **LOD:** limit of detection.

## Discussion

Multiple protein engineering strategies have been investigated to generate cross-protective or epitope-based antigens against RSV F or hMPV F. Previous attempts at grafted a single antigenic site on RSV F or hMPV F from one to another induced cross-neutralizing antibodies but very limited protection against the heterologous virus challenge, likely due to the epitopes on the backbone still dominate the immune responses (39). RHMS-1 contains multiple immunodominant epitopes of both RSV F and hMPV F in relatively equal propotions, including at least three RSV F-specific and three MPV F-specific antigenic sites. Therefore, a more balanced immune responses against both RSV F and hMPV F would be expected, and it is less likely to drive escape mutations focused on a single epitope. After prime and boost, RHMS-1 induced comparable levels of hMPV F/RSV F-specific serum IgG titers. Although the serum neutralization against RSV is not as potent as that against hMPV, RHMS-1 immunization completely protected the mice from both RSV and hMPV challenges, suggesting RHMS-1 is a promising antigen that can be used as a vaccine to induce cross-neutralizing and cross-protecting antibodies against RSV and hMPV. These findings also suggest additional optimization of antigen and adjuvants are warranted to optimize the elicited neutralizing antibody responses.

Our ELISA screening data showed that the human subjects tested had pre-existing immunity against both RSV F and hMPV F. Interestingly, subjects had an overall higher frequency of RSV F-specific B cells than hMPV F-specific B cells, which is likely due to the higher prevalence of RSV than hMPV. Since initial exposures to an antigen can influence subsequent immune responses against similar antigens, termed original antigenic sin (49), pre-existing immunity to RSV or hMPV may affect the efficacy of RHMS-1 or other RSV and hMPV F-based vaccine candidates. To address this problem, future studies will be needed to test RHMS-1 in animal models that have exposed to RSV and hMPV. Furthermore, despite encouraging results obtained from the RHMS-1 immunization/challenge in this study, it is still necessary to unserstand if RHMS-1 works better than a mixture of RSV F and hMPV F in animal models.

Previous studies have shown that mice immunized with either pre-fusion RSV F or post-fusion hMPV F did not induce significant cross-neutralization antibodies (17). Similar results were observed in this study with pre-fusion RSV F and monomeric hMPV F. The serum of 9 out of 16 mice immunized with RSV F + AddaS03 showed little binding to monomeric hMPV F just above the detection limit, while all of the monomeric hMPV F + AddaS03 immunized mice serum had moderate binding to RSV F. However, the serum of both groups failed to cross-neutralize the viruses *in vitro* **(Figure 6B-E)**. Interestingly, the lung virus counts in RSV F immunized/hMPV challenged and hMPV F immunized/RSV challenged groups are reduced ∼10 fold compared to mock immunized/hMPV challenged and mock immunized/RSV challenged groups respectively **(Figure 6F)**, indicating poorly and non-neutralizing, cross-reactive antibodies could play a role in limiting virus replication in the lungs of mice, possibly through Fc-mediated effector functions that need to be further characterized. In addition, future animal studies in more permissive animal models such as cotton rats and African Green Monkeys need to be performed to further evaluate the efficacy and safety of RHMS-1.

Although the RHMS-1 construct appears to be a pre-fusion trimer by negative-stain electron microscopy, the stability of this protein requires further evaluation, and could likely be optimized to improve both stability and antigenicity using computational protein design tools. For example, by using interprotomer disulfides (IP-DSs) that linking protomers of the hMPV F trimer, both stabilized pre-fusion and post-fusion F proteins elicited significantly higher neutralizing responses than the hMPV F proteins without IP-DSs (50). Such a strategy could also be applied to stabilize the RHMS-1 construct. In addition, assessment of the fusogenicity of RHMS-1 will need to be assessed to determine if incorporation into live-attenuated vaccine platforms is warranted.

In summary, we generated and evaluated a chimeric RSV F and hMPV F protein, RHMS-1. To our knowledge, this is the first immunogen that elicits potent and balanced protective immune responses against both RSV and hMPV. Moreover, RHMS-1 can be readily applied to both traditional and novel vaccine delivery platforms like viral vectors, VLPs, nanoparticles, and mRNA. Further optimization of RHMS-1 could lead to a safe and effective universal Pneumovirus vaccine.

## Acknowledgement

These studies were supported by National Institutes of Health grants 1R01AI143865 (J.J.M.) and 1K01OD026569 (J.J.M.). The funders had no role in study design, data collection and analysis, the decision to publish, or preparation of the manuscript. Molecular graphics and analyses performed with UCSF ChimeraX, developed by the Resource for Biocomputing, Visualization, and Informatics at the University of California, San Francisco, with support from National Institutes of Health R01-GM129325 and the Office of Cyber Infrastructure and Computational Biology, National Institute of Allergy and Infectious Diseases.

## Author contribution

J.H. and R.M. conducted the experiments. J.H., R.M., and J.J.M. conducted the data analysis and wrote and revised the paper.

## Data availability

The authors declare that all data supporting the findings of this study are available within the paper.

